# Genetic biosensor for optimizing double-stranded RNA production by bacterial symbionts

**DOI:** 10.64898/2026.01.18.700149

**Authors:** Lucio Navarro-Escalante, Anthony J. VanDieren, Jeffrey E. Barrick

**Author notes:** Correspondence: L. N.-E. and J.E.B.

## Abstract

There is growing interest in engineering animal and plant microbiomes to deliver double-stranded RNA (dsRNA) for RNA interference (RNAi) applications. We developed a genetically encoded biosensor that uses bimolecular fluorescence complementation to monitor dsRNA levels within bacterial cells to accelerate the symbiont-mediated RNAi design–build–test cycle. We validated performance of the sensor in *Escherichia coli* and demonstrated enhanced dsRNA accumulation in engineered strains of the aphid symbiont *Serratia symbiotica*.

## Main Text

Several methods have been developed to detect and visualize double-stranded RNA (dsRNA) in cells and tissues, including immunofluorescence assays and the use of dsRNA-binding domains fused to fluorescent proteins^1–3^. These tools have been primarily designed to detect viral dsRNA molecules in eukaryotic cells for studying and diagnosing infections of mammalian and plant viruses^4,5^. They are not suitable for the in vivo quantification of dsRNA molecules in prokaryotic cells. Here, we describe a genetically encoded dsRNA biosensor that provides a fluorescent readout of overall dsRNA levels in living bacterial cells.

Our dsRNA-sensing system is based on bimolecular fluorescence complementation (BiFC). Non-fluorescent fragments of a split mVenus yellow fluorescent protein (YFP) reporter are expressed as fusion proteins containing dsRNA-binding domains such that fluorescence is reconstituted when they associate with the same dsRNA molecule (**Fig. 1a**). We tested dsRNA-binding domains (RBDs) from two well-characterized viral proteins, B2 from flock house virus and NS1 from influenza A virus. These proteins bind dsRNA in a sequence-independent manner and have been used to detect and visualize viral dsRNA in plant tissues^2,3,6–8^. We constructed four plasmids (pDS-BiFC1 through pDS-BiFC4), encoding different RBD fusions to the split mVenus N-terminal and C-terminal fragments expressed from T7 promoters (**Fig. 1b**).

**Fig. 1.**
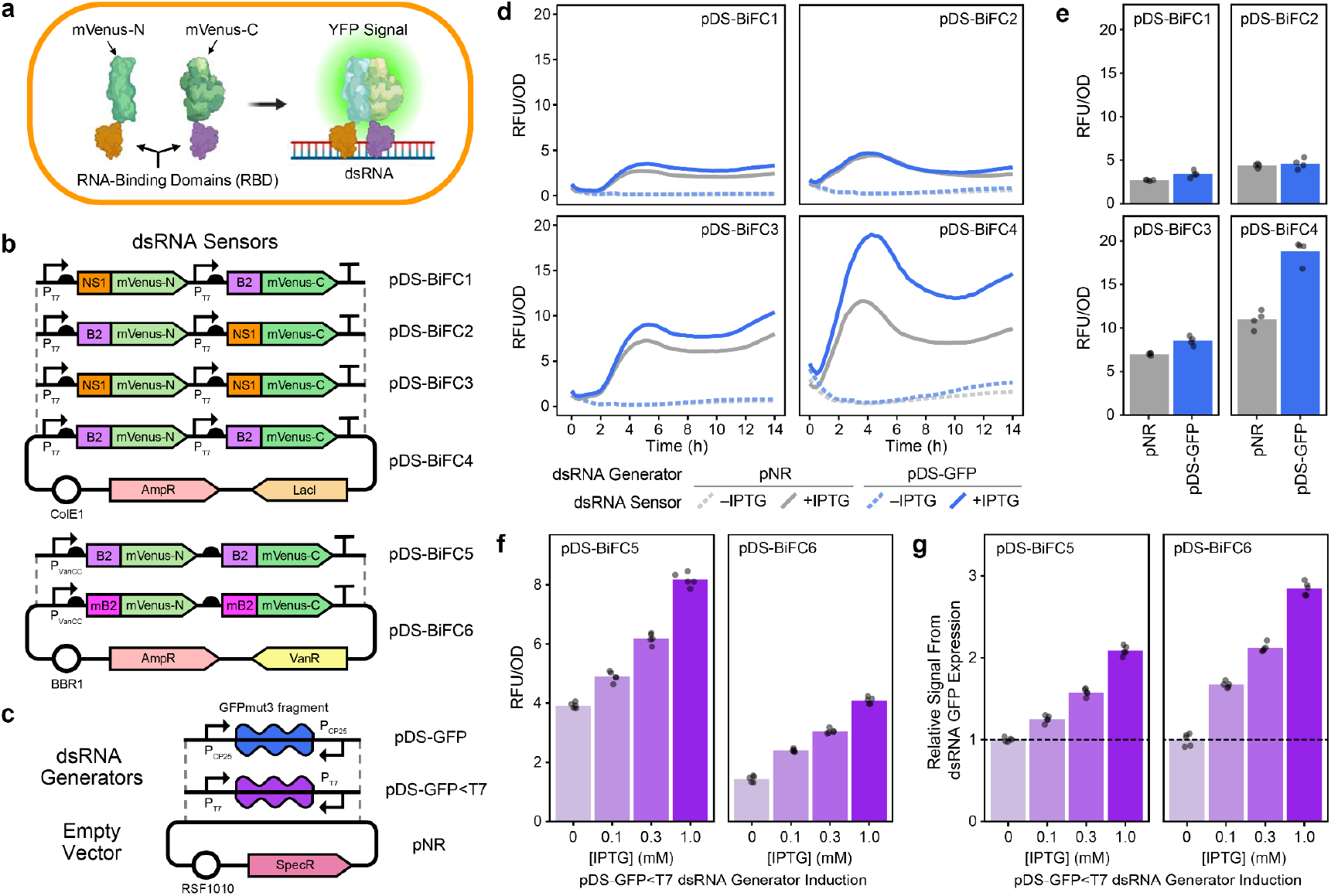
dsRNA biosensor optimization in *E. coli*. **a**, Bimolecular fluorescence complementation yields a yellow fluorescence protein (YFP) signal when dsRNA binding proteins bring together the two halves of mVenus. **b**, Plasmid maps for dsRNA sensor designs. **c**, Plasmid maps for dsRNA generators used to test sensor performance and the empty vector control. **d**, Time courses of YFP relative fluorescence units (RFU) per optical density at 600 nm (OD600) in *E. coli* cultures containing initial IPTG-inducible dsRNA sensor designs. Curves show the mean of four replicates. **c**, Values from time courses in **d** at 4.5 h. **f**, YFP RFU/OD600 signal for vanillate-inducible sensor designs 6.5 h into a time course after adding vanillate (Van) and inducing dsRNA production with different concentrations of IPTG. **g**, Values in **f** normalized to the uninduced (0mM IPTG) dsRNA generator control.

To test whether these constructs could detect dsRNA expression, we transformed them into *Escherichia coli* HT115 (DE3), a strain commonly used for dsRNA production due to its IPTG-inducible T7 RNA polymerase and lack of RNase III activity^9^. These strains were further co-transformed with either an empty vector (pNR) or a dsRNA generator plasmid that constitutively expresses a 717-bp dsRNA fragment from GFPmut3 gene (pDS-GFP)^10^ (**Fig. 1c**). Fluorescence measurements after IPTG induction of the sensors (**Fig. 1d**,**e**) revealed that the heterotypic designs, in which NS1 and B2 RBDs were paired, yielded little to no additional YFP fluorescence in strains with the dsRNA generator plasmid. In contrast, the homotypic designs, particularly the B2-B2 RBD configuration, exhibited much higher YFP fluorescence. Therefore, we selected the pDS-BiFC4 design for further optimization.

Because our goal was to create a broadly applicable tool for dsRNA detection in bacteria, including non-model species such as insect symbionts, we transferred the pDS-BiFC4 design into a broad-host-range plasmid backbone (BBR1 origin) and replaced the T7 promoter with a vanillate-inducible promoter system^11^, generating pDS-BiFC5 (**Fig. 1b**). Because B2 can dimerize, possibly leading to dsRNA-independent reconstitution of the mVenus reporter, we also introduced an amino acid substitution known to eliminate B2 dimerization^12^ in an alternative design, pDS-BiFC6 (**Fig. 1e**). We again tested sensor performance in *E. coli* HT115 (DE3) but co-transformed this time with a dsRNA generator plasmid that expresses the dsRNA fragment from GFPmut3 under control of an IPTG-inducible T7 promoter (pDS-GFP<T7). Both pDS-BiFC5 and pDS-BiFC6 responded with increased fluorescence as higher concentrations of IPTG were added to trigger more dsRNA expression (**Fig. 1f**). pDS-BiFC6 consistently showed a higher relative dsRNA-specific signal than pDS-BiFC5 (**Fig. 1g**). On this basis, we selected pDS-BiFC6 as the optimized version of our dsRNA sensor.

We next examined whether our broad-host-range dsRNA-sensor plasmid functioned in the culturable aphid symbiont *Serratia symbiotica* CWBI-2.3^T^ (henceforth CWBI)^13–15^ (**Fig. 2a**). There was no detectable difference in fluorescence between *S. symbiotica* strains containing the pDS-BiFC6 sensor plasmid after they were co-transformed with either a pDS-Def1 dsRNA generator plasmid (**Fig. 2b**), which expresses a 184-bp fragment of the honey bee (*Apis mellifera*) defensin-1 gene^16^, or the pNR empty vector control, suggesting that our vanillate-induction system does not function in this insect symbiont. To overcome this limitation, we constructed pDS-BiFC7, which uses the pBAD arabinose-inducible promoter system to control dsRNA sensor expression (**Fig. 2c**).

**Fig. 2.**
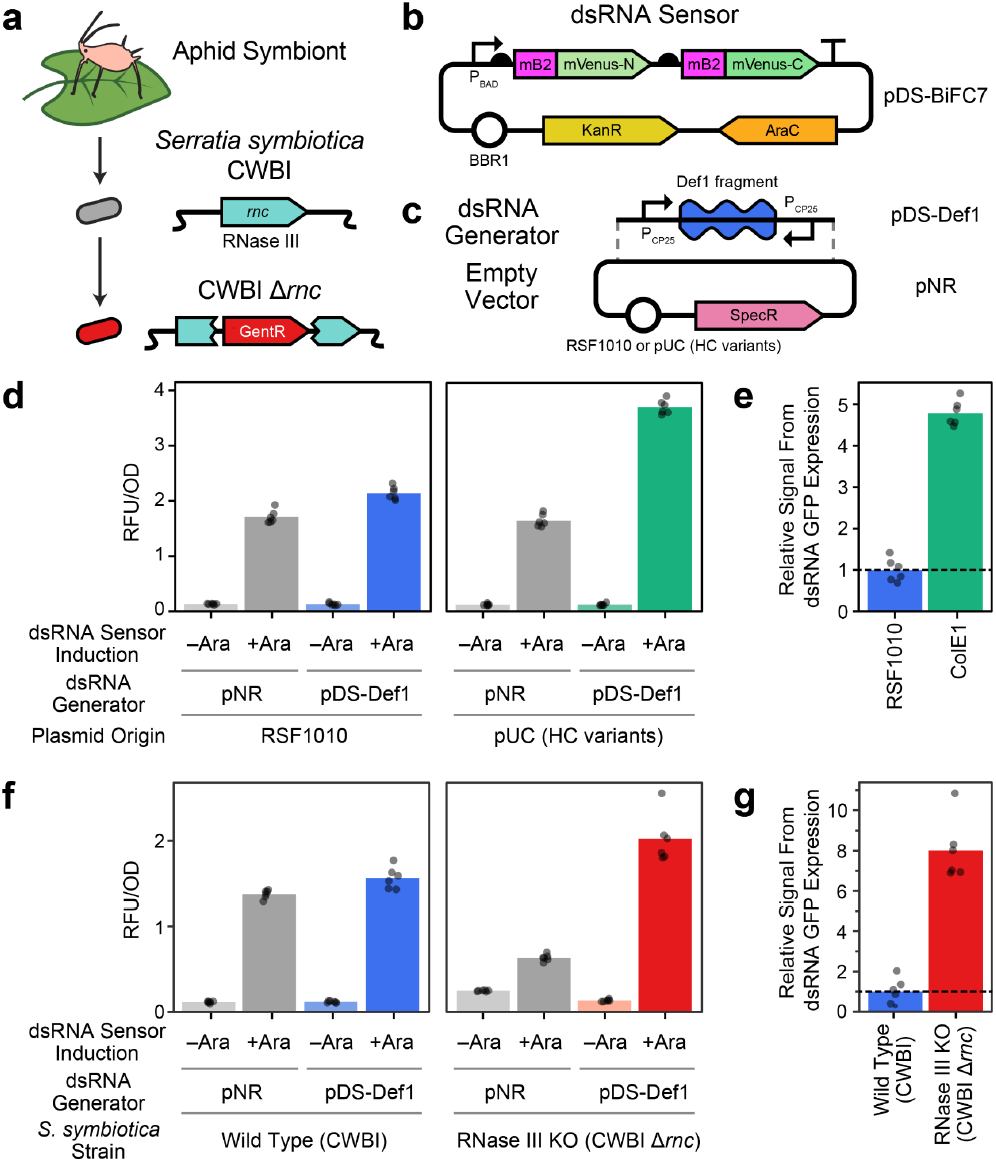
Improved dsRNA accumulation in engineered *Serratia symbiotica*. **a**, The bacterium *S. symbiotica* CWBI is a culturable aphid symbiont. We engineered the RNase III knockout CWBI Δ*rnc* strain by disrupting this gene with a gentamicin resistance cassette. **b**, Plasmid map for the arabinose-inducible dsRNA sensor design used in *S. symbiotica*. **c**, Plasmid maps for dsRNA generators and corresponding empty vector controls used in *S. symbiotica*. **d**, Comparison of dsRNA sensor signal after 24 h from the dsRNA generator and empty vector control plasmids with the medium-copy RSF1010 origin of replication relative to high-copy (HC) variants with the pUC origin with and without arabinose (Ara) induction of the sensor. **e**, Data in **d** with signal from empty vector control background subtracted and then normalized to the RSF1010 plasmid value. **f**, Comparison of dsRNA sensor signal in *S. symbiotica* CWBI and CWBI Δ*rnc* strains with the RSF1010 dsRNA generator plasmid or empty vector control. **g**, Data in **f** with signal from empty vector control background subtracted and then normalized to the CWBI strain value.

Using pDS-BiFC7, we compared constitutive dsRNA expression in *S. symbiotica* CWBI from two different plasmid backbones: one with a medium-copy RSF1010 origin and one with a high-copy pUC origin (**Fig. 2b**). Upon sensor induction with arabinose, both plasmids produced significant increases in fluorescence compared with their respective empty vector controls (Welch”s *t*-test, *p* < 0.0001) (**Fig. 2d**). pDS-BiFC7 generated ∼4.8-fold higher fluorescence for the pUC plasmid compared with the RSF1010 plasmid, indicating that using this high-copy plasmid greatly increased dsRNA accumulation (**Fig. 2e**).

To further test the utility of our sensor for comparing symbionts engineered for increased dsRNA production, we generated *S. symbiotica* CWBI Δ*rnc*, a strain with an antibiotic resistance cassette inserted so that it disrupts the RNase III gene (**Fig. 2a**). Attempts to introduce the high-copy pUC version of pDS-Def1 into CWBI Δ*rnc* repeatedly yielded colonies with truncated plasmids missing the dsRNA expression cassette, suggesting that dsRNA accumulation in the absence of RNase III can become toxic. Therefore, we measured dsRNA production in this strain using the lower copy RSF1010 version of pDS-Def1. When co-transformed with pDS-BiFC7, the mutant CWBI Δ*rnc* strain showed ∼8-fold higher dsRNA signal compared with the wild-type CWBI strain, as measured relative to the pNR control (**Fig. 2f**,**g**). These results confirm that deletion of RNase III enhances dsRNA accumulation in *S. symbiotica* CWBI.

In summary, we developed a genetically encoded dsRNA biosensor for bacteria. By fusing viral dsRNA-binding domains to split mVenus and expressing these proteins from plasmids with different origins of replication and under control of different inducible promoter systems, we showed that this sensor design is versatile and robust. We used the dsRNA sensor to demonstrate that newly engineered variants of the aphid symbiont *S. symbiotica* CWBI accumulate more of a heterologously expressed dsRNA than strains and plasmids used in prior tests of symbiont-mediated RNAi in this insect^13^. These sensors can be used to develop more efficient bacterial systems for large-scale dsRNA manufacturing^17–19^ and support growing interest in symbiont-mediated RNAi applications^20–22^.

## Methods

### Bacteria strains and media

*E. coli* DH5α was used for all plasmid cloning and maintenance. *E. coli* HT115 (DE3) was used for dsRNA sensing assays. *E. coli* was cultured in the Miller formulation of lysogeny broth (LB) at 37°C. *Serratia symbiotica* CWBI-2.3T was cultured in tryptic soy broth (TSB) or agar (TSA) at 25°C. Media were supplemented with antibiotics at the following concentrations for each resistance cassette: 100 µg/mL carbenicillin (AmpR), 60 µg/mL spectinomycin (SpecR), 50 µg/mL kanamycin (KanR), and 20 µg/mL gentamicin (GentR). Inducers were added at these concentrations unless otherwise indicated: 0.2 mM isopropyl β-D-1-thiogalactopyranoside (IPTG), 0.2 mM vanillate (Van), and 0.2% w/v arabinose (Ara). **Plasmid construction**. Plasmids used and created in this work are listed in **Supplementary Table 1**. Sequences of oligonucleotides and DNA fragments are provided in **Supplementary Table 2**. Plasmids pDS-BiFC1 to pDS-BiFC4 were constructed by using circular polymerase extension cloning (CPEC) (Quan and Tian 2009) to insert DNA fragments encoding NS1 and B2 RNA-binding domains synthesized with appropriate overhangs into pET-BiFC (Eastwood et al. 2017). Plasmid pDS-BiFC5 was created by combining Type 3a B2-mVenus-C and Type 3b B2-mVenus-C DNA fragments, pBTK1075 (Type 2 P_Van_ promoter), pBTK1076 (Type 4 VanR repressor), and pBTK1063 (Type 5-1 pBBR1 dropout) using BsaI Golden Gate assembly (GGA) according to the YTK/BTK standard^23,24^. pDS-BiFC6 was created by site-directed mutagenesis of pDS-BiFC5 using a 2-step CPEC approach. To create pDS-BiFC7, we first cloned the sensor from pDS-BiFC6 into a pBAD plasmid. Then, the P_BAD_ promoter, sensor, and AraC transcriptional unit (amplified as a Type 2-4 part) were cloned into pBTK1116 (Type 5-1 pBBR1 backbone) using BsaI GGA. pNR-HC and pDS-Def1-HC were created by BsaI GGA of the insert from either pNR or pDS-Def1 dsRNA expression plasmid (amplified as a Type 2-4 part) into pBTK1213 (Type 5-1 pUC dropout).

### dsRNA sensing assays in *E. coli*

Overnight cultures of *E. coli* containing a dsRNA sensor plasmid and a dsRNA generator plasmid or the empty vector control were diluted to an optical density at 600 nm (OD600) of 0.2. Inducers were added for the sensor and, if applicable, the dsRNA generator as specified. Then, 200 µl aliquots of these cultures were distributed into wells of a black-walled, glass-bottomed 96-well plate that also included media blank controls. Microplate incubation and OD600 and YFP fluorescence readings were performed in a Tecan Infinite Pro M200 Plate Reader using 500 nm for excitation and 538 nm for emission. Cultures were incubated for a total of 14 h with readings taken every 15 min over the time course with 13 min of orbital shaking between each set of measurements.

### Construction of *S. symbiotica* CWBI Δ*rnc*

RNase III was disrupted by inserting a CRISPR-associated minitransposon into the *rnc* gene using previously described methods^25^. In brief, an *rnc* guide RNA sequence (Supplementary Table 2) and a GentR cassette were cloned into the UltraCAST vector via Golden Gate assembly. The assembled vector was conjugated from *E. coli* MFDpir into *S. symbiotica* CWBI. A GentR colony was selected, and its genome was Illumina sequenced to confirm knockout of the *rnc* gene.

### dsRNA sensing assays in *S. symbiotica*

dsRNA sensor and generator plasmids were electroporated into *S. symbiotica* as previously described^15^, except using a voltage of 1.5 kV. After growth to saturation, we diluted cultures of strains being tested to an OD600 of 0.2 and added arabinose for sensor induction as specified. Then, 300 µl aliquots of these cultures were distributed into wells of a black-walled, glass-bottomed 96-well plate that also included media blank controls. OD600 and YFP fluorescence were measured as for *E. coli*, except *S. symbiotica* time courses lasted a total of 24 h.

## Data availability

Whole-genome sequencing data for the *S. symbiotica* CWBI *Δrnc* strain is publicly available from the NCBI Sequence Read Archive (PRJNA1347086). Plasmids are being deposited in Addgene (https://www.addgene.org/plasmids/articles/28264021/). All other data supporting the findings of this study are available within the paper and its Supplementary Information.

## Author contributions

Conceptualization: L. N.-E.; Investigation: L. N.-E. and A.J.V.; Writing - Original Draft: L.N.-E. and A.J.V.; Writing - Review & Editing: L. N.-E., A.J.V, and J.E.B. Visualization: L.N.-E. and J.E.B.; Funding acquisition: J.E.B.

## Acknowledgments

This work was funded by the U.S. National Science Foundation (IOS-2103208) and the U.S. Army Research Office (W911NF-20-1-0195).

---

**Supplementary Table 1** | Plasmids used in this study.

**Supplementary Table 2** | Primers, oligos, and gene fragments used in this study.

**Source Data** | Raw and processed underlying data for figures.

